# Tell me who your neighbors are: The role of spatial location and tree species identity in determining the ectomycorrhizal community composition of saplings and mature trees in a mixed conifer forest

**DOI:** 10.1101/2022.10.27.514003

**Authors:** Stav Livne-Luzon, Mor Avidar, Lior Herol, Ido Rog, Tamir Klein, Hagai Shemesh

## Abstract

1. The mutualistic interaction between trees and ectomycorrhizal fungi (EMF) can have a major effect on forest dynamics and specifically in seedling establishment. Both intrinsic (i.e., identity of the sapling) and extrinsic (i.e., the identity of mature trees in the vicinity of the sapling) factors can affect the EMF community composition of young saplings.
2. Here, we compared the EMF community composition associated with the roots of young saplings and mature trees of two co-habiting Pinaceae: *Pinus halepensis* and *Cedrus deodara* growing together in a planted forest plot, using fungal ITS metabarcoding.
3. We found that the differences between the two sapling groups were mostly attributed to changes in the relative abundance of specific fungal species. Moreover, we found that physical proximity to a specific host species had a significant effect on the community composition of young saplings. However, while no significant differences in sapling size were apparent, the sapling shoot structure was affected by the identity of the nearest mature tree and its unique EMF community composition.
4. *Synthesis*: These results suggest that the dynamics of the EMF community are greatly determined by extrinsic factors such as the small-scale distribution of mature trees in the forest, with possible cascading effects on the development of young trees.

## Introduction

The world’s forests are one of the largest reservoirs of carbon and as such, they play a crucial role in the global carbon cycle. However, recent changes in climate are expected to affect forests throughout the globe, potentially leading to a positive feedback that could result in the deterioration of forests and accelerated global warming (Adams et al., 2009, Kurz et al., 2008). Seedling establishment is crucial for forest persistence and often restricts population growth due to high mortality during the early growth stages (Fenner, 1987). Such mortality can arise from both abiotic conditions, such as drought (Engelbrecht et al., 2005) and extreme temperatures (Schaberg et al., 2008), as well as biotic interactions such as inter-plant competition and herbivory (Meiners and Handel, 2000). However, biotic interactions are not necessarily detrimental and can also have positive effects on seedling establishment.

For many tree species, successful seedling establishment depends on early association with soil symbionts (Dickie and Reich, 2005, Van Der Heijden et al., 2016). Plant interactions with mycorrhizal fungi are one of the most common and ecologically important mutualisms in nature (Smith and Read, 1997). This is especially true in temperate and boreal forests, where many of the dominant tree species (e.g., pine, oak, spruce, larch & beech) are obligatorily associated with ectomycorrhizal fungi (EMF). This dependence can have far reaching consequences on vegetation’s successional trajectories (Reynolds et al., 2003, Hartnett and Wilson, 2002), including the ability of forests to expand (Frank et al., 2009) and regenerate after large scale disturbances such as fire (Glassman et al., 2016) and clearcutting (Grove et al., 2019).

Due to the pivotal role of seedling establishment in forest regeneration, it is critical to understand the mechanisms governing the dynamics of the EMF community on the roots of tree seedlings and how this community changes as the host tree ages. Various intrinsic and extrinsic factors can shape the EMF community composition. Some intrinsic factors relate to the biological contexts of the hosts themselves such as host identity (Ishida et al., 2007) (Tedersoo et al., 2016, Otsing et al., 2021, Bogar and Kennedy, 2013) and age (Jonsson et al., 1999) while other extrinsic factors relate directly to the abiotic characteristics of the habitat such as soil properties (Erlandson et al., 2018) and moisture (Gehring et al., 2006). Nevertheless, EMF colonization takes place as roots encounter spores or existing hyphae, and therefore extrinsic biotic factors such as the spatial arrangement of the inoculum sources in relation to the seedling’s roots might play a major role in determining seedling establishment (Livne-Luzon et al., 2017, Livne-Luzon et al., 2021). Therefore, the identity and spatial arrangement of neighboring mature trees should affect the spatial distribution and the identity of the EMF inoculum seedlings encounter.

The encounter of plant roots and EMF inoculum is not completely random and certain levels of specificity are known to occur in host-fungal interactions (Molina and Horton, 2015). Such host preference has been suggested to explain the positive relation between plant diversity and symbiont diversity (Kernaghan et al., 2003). The interaction among members of the Pinaceae and fungi of the *Suilloid* group are a good example for such high level of specificity in host-mycorrhizal interactions (Horton and Bruns, 1998, Dahlberg and Finlay, 1999). Such phylogenetic associations are not temporally constant and can change as the host ages. For example, while fungi from the *Suilloid* group are common on the roots of seedlings and young trees in the Pinaceae, they are rare on mature trees which often associate with fungi characteristic of late succession such as *Russula* and *Sebacina* species (Rog et al., 2020). Furthermore, in addition to the changes in community structure, forest maturation is often associated with an increase in EMF richness and diversity (Odriozola et al., 2020).

Both *Pinus halepensis* and *Cedrus deodara* belong to the Pinaceae and obligately interact with ectomycorrhizal fungi (Molina et al., 1992). *Pinus halepensis* is native to Israel and has a widespread distribution across all of Israel. While other *Cedrus* relatives (i.e., *Cedrus libani*) have a long history of inhabiting specific sites in Israel, it has vanished from the pollen record ~ 900 ybp (Bar-On et al., 2022). In contrast, *Cedrus deodara* is a foreign species with a patchy distribution limited to specific areas where it has been planted. Nevertheless, both species exhibit very low natural regeneration within dense forests (where their establishment is thought to be hindered by competition). These two co-habiting species provide an opportunity to examine some of the extrinsic and intrinsic biotic factors which shape the EMF communities of young trees. More specifically, we test whether similarity in EMF community composition is dominated by host identity, host age, or mature neighbor identity. We hypothesized that the saplings of the two tree species should be distinct in their ectomycorrhizal community composition, while mature trees will be more similar. Additionally, because the spatial distribution of mature trees should be linked to inoculum potential, we expected it to affect the EMF community composition of young saplings.

## Materials and methods

### Study site

The study was conducted in northern Israel, in part of the nature reserve of Mt. Meron in the upper galilee (N 35°24’11, E”05’33°00; elevation 850 m above sea level, aspect: north-west, slope: 7-15°, average annual precipitation 900 mm). The study site is characterized as Mediterranean woodland on Terra rosa soil and is dominated by *Quercus calliprinos* and *Pistacia palaestina*. Additionally, ~70 years ago both *Pinus halepensis* and *Cedrus deodara* were planted in the area and have both been demonstrating nearly zero regeneration within the plot. However, a fire that occurred ~7 years prior to our study (2014) created a clearing that facilitated the regeneration of a few dozen seedlings of pines and cedar (Fig. S1).

### Tree species

Several characteristics could explain the post-fire regeneration of *Pinus halepensis* within the experimental plot. First, it reaches reproductive maturity at a relatively young age and is a prolific producer of serotinous, wind-dispersed seeds. Second, it is a highly drought resistant tree (Klein et al., 2011) and is habituated in one of the driest and most flammable of all areas in the Mediterranean Basin. Pines are known to associate with various taxa of EMF (e.g., *Suillus, Rhizopogon, Amanita, Pisolithus, Lactarius* and *Tuber*). On the contrary, *Cedrus deodara* is considered less resistant to drought and calcareous soils and in Israel it was mostly planted (and survived) in areas where the average precipitation exceeds 600 mm/year. Like other cedars, it does not seem to have any fire-prone adaptations and its increased regeneration in the post-fire site described above, could be merely related to reduced competition with annual plants and grasses. Little is known of the EMF communities of *Cedrus deodara*, especially out of its native range, but preliminary assessment of its EMF community in the Himalayas (Pande et al., 2004) showed association with various taxa such as: *Boletus*, *Russula* and *Amanita*. Other studies demonstrated a potential association with *Rhizopogon* (Thakur et al., 2021) and *Scleroderma* (Itoo and Reshi, 2014) under nursery conditions. The rooting systems of the mature *Pinus halepensis* and *Cedrus deodara* differ greatly. In *Pinus halepensis* the root system is quite significant, with a shoot:root ratio of 1:10 (Grünzweig et al., 2007). However, the overall diameter of the rooting system is usually smaller than the diameter of the canopy and most of the roots are found in the shallow soil layers (1-2m) with only several roots reaching to deeper layers of up to 6m (Rog et al., 2021). The *Cedrus deodara* rooting system appears to be less substantial, with a shoot:root ratio of 1:4 and roots which were documented to reach up to 2.4 m (in nineteen year old trees; Wani et al., 2014). In this current study we examined the ectomycorrhizal communities of four groups: mature pines, mature cedars, young pines and young cedars (Fig. S1).

### Young sapling sampling

Prior to sampling, the upper soil and litter layer (~1 cm depth) around each sapling was removed, then each sapling was carefully removed by digging gently around its rooting system, leaving as much of the roots as intact as possible. Then, fine roots which had apparent mycorrhizal root tips were collected into a moist Ziplock bag and kept cold (4 °C) until further inspection in the lab. For each sapling we measured the stem diameter (cm) using a digital caliper (Signet, Taichung city, Taiwan), the sapling shoot length (in centimeters from soil surface to uppermost shoot tip) and the number of side branches. We also documented, for each sapling, its distance (cm) from the nearest mature tree and the identity of that nearest neighbor.

### Mature tree sampling

Mature tree sampling followed the methods of Rog et al. (2020). Briefly, we chose five pairs of adjacent mature cedars and pines (with an average distance of 4.8 m). Similar to the sapling sampling, we cleared the upper soil and litter layer around each tree. Lateral fine roots (4-5 for each individual tree) were sampled at ~50 cm distance from stems they belong to, at the top soil, about 10 cm soil depth; tree species was controlled by following roots to the stems they emanated from (and by DNA sequence analysis of roots; detailed below). For each root we recorded the distance and orientation to the source tree. Sampled roots were stored in an ice box and washed from attached soil particles under tap water within 24 hours. Washed fine roots were placed in Petri dishes with a thin layer of unchlorinated tap water and stored at 4 °C for 24 hours.

### Fungal root-tip collection

Roots of each individual sapling and mature tree roots were carefully washed in a sieve (2 mm) and all colonized root tips were removed using sterilized forceps, inserted into a 1.5 ml Eppendorf tube added with 300 μl CTAB +PVPP 2% buffer, and immediately stored in a 20°C freezer until DNA extraction. To avoid cross-contamination between samples, all tools were sterilized using ethanol (70%).

### Molecular identification of fungal species

DNA extractions followed the methods of Livne-Luzon et al., (2017). Briefly, frozen root tips were bead beaten (2 × 30 seconds at 4000 rounds per minute), and DNA was extracted from each root tip sample following a modified version of the QIAGEN (Valencia, CA, USA) DNAeasy Blood and Tissue Kit. Barcoded amplicon sequencing of the fungal ITS2 region was performed on a MiSeq platform (Illumina, San Diego, CA, USA). A two-step protocol for library preparation was performed according to Straussman lab (Nejman et al., 2020) with several modifications described in detail in Avital et al., 2022; First PCR reactions were performed using KAPA HiFi HotStart ReadyMix DNA polymerase (Hoffmann-La Roch, Basel, Switzerland) in 50 μl reaction volumes with 5 μl DNA extract, and 1 μl of every primer 5.8S-Fun (5’- AACTTTYRRCAAYGGATCWCT) (Tylor et al., 2016) and RD2-ITS4Fun (5’AGACGTGTGCTCTTCCGATCT-AGCCTCCGCTTATTGATATGCTTAART). The reverse primer consisted of the ITS4-Fun primer (Tylor et al., 2016) with the linker adapter RD2. PCR reactions were performed as follows: initial 2 min at 98 °C followed by 35 cycles of 10 s 98 °C, 15 s 55 °C, and 35 s 72 °C final cycle with 5 min 72 °C. Second PCR reactions were performed using the same DNA polymerase in 50 μl reaction volumes with 1/10 (5 μl) of the first PCR reaction, and 1 μl of every primer P5-rd1-5.8S-Fun (5’- AATGATACGGCGACCACCGAGATCT-ACACTCTTTCCCTACACGACGCTCTTCCGATCT-AACTTTYRRCAAYGGATCWCT) and RD2-Barcode (5’ AGACGTGTGCTCTTCCGATCT-BARCODE). The forward primer consisted of the adaptor p5, the linker RD1, and the primer 5.8S-Fun. The reverse primer consisted of the adapter RD2 and the individual barcode. PCR reactions were performed similarly to the first PCR but with only 6 cycles. PCRs were cleaned using Qiaquick PCR purification kit (Qiagen, Hilden, Germany), quantified fluorescently with the Qubit dsDNA HS kit (Life Technologies Inc., Gaithersburg, MD, USA). Libraries were quality checked for concentration and amplicon size using the Agilent 2100 Bioanalyzer (Agilent Technologies, Santa Clara, CA, USA) and size selected with AMPure magnetic beads (Beckman Coulter Inc., Brea, CA, USA). We sequenced all the samples in one amplicon using Illumina MiSeq technology with 300 bp paired-end reads (PE300_V3) in the Grand Israel National Center for Personalized Medicine (Weizmann institute of science, Rehovot, Israel).

### Validation of tree species identity

Mature tree roots were verified for species identity by the tree ITS-2 region. We amplified the tree ITS-2 region and verified our assessment of tree species identity. PCR reactions were performed using 1.25 units Dream Taq DNA polymerase (Thermo Scientific) in 25 μl reaction volumes with 1 μl DNA extract, and 1 μl of every primer ITS2-S2L (5’-ATGCGATACTTGGTGTGAAT) and ITS2-S3R (5’-GACGCTTCTCCAGACTACAAT) (Yao H., et al 2010). PCR reactions were performed as follows: initial 1 min at 94 °C followed by 35 cycles of 30 s 94 °C, 30 s 56 °C, and 1 min 72 °C final cycle with 8 min 72 °C. Six μl of the PCR products were run on 1.5% agarose gel and PCR reaction samples with one clear band were purified by Exo-1 and rSAP and Sanger sequenced by Biological Services Department (Weizmann Institute of Science, Rehovot, Israel), uni-directionally with primer ITS2-S3R. The ITS2 sequences were blasted manually in the NCBI dataset. Only samples with clear tree species identity; >95% identity high query cover were further analyzed. Overall, although carefully picked, only 28 roots out of the 47 roots sampled were correctly classified, one sample was classified as *Quercus*, one as *Pistacia*, for an additional seventeen samples we could not verify the sampled tree. Further analysis was performed based on the verified identity of the hosts.

### Bioinformatics

We used R (R Core Team, 2018, version 4.0.3) and the R-Studio IDE for bioinformatics and statistical analysis. Raw sequences were demultiplexed and adapters together with barcodes were removed for 84 root samples (37 saplings, 47 mature of 10 individual trees). The sequences were analyzed using the amplicon sequencing dada2 package v. 1.7.9 in R (Callahan et al., 2016). In summary, sequences were quality-filtered and trimmed. We only used sequences longer than 50 bases with a mean number of expected errors below 2 (maxN = 0, maxEE = c(2,5) minLen = 50 truncQ = 2). Paired-end sequences were merged using the MergePairs function. We then applied a dereplication procedure on each sample independently, using derepFastq function. Finally, all files were combined in one single Fasta file to obtain a single amplicon sequence variant (ASV) data file. We removed singletons (minuniquesize = 2) and de novo chimera sequences using removeBimeraDenovo function against the reference database (UNITE/UCHIME reference datasets v.7.2). Sequences were then clustered, and taxonomic assignment (id = 0.98) was done against the UNITE database. Non-fungal ASVs were removed. FUNguild was then used to parse ASVs into ecological guilds (Nguyen et al. 2016) we then filtered the table to include only fungal taxa which were scored as highly probable ectomycorrhizal by the FUNguild algorithm. We had 3 negative control samples: one with ultrapure water, one with all PCR reagents and one having only the second PCR reagents. The negative control samples had several ASVs with low reads (~400 total). To avoid contamination, we summed all ASVs that were found on them and removed these from each of the other samples. The resulting ASV table was agglomerated at the species level and was narrowed to include only the most abundant species i.e., species which had at least 100 reads across all samples. Qualitative similar results were obtained while agglomerating the sequence data to the genus level.

### Statistical analysis

#### Fungal community richness

Fungal richness (the number of “species” with read abundance greater than zero in each sample) was calculated at two levels: 1) at the raw ASV level (prior to any filtering for EMF species or agglomeration of the data). 2) After agglomeration of the data to the EMF species level and filtering the table to include only species with more than 100 reads in all samples. On both levels, we used a general linear model using a fully factorial design to account for differences in richness. The following explanatory variables were included as fixed factors: the identity of the tree (cedar/pine), the host age (mature/saplings) as well as their interaction.

#### Fungal community composition

To illustrate the main axes discriminating between treatments (host identity and host age), a permutational MANOVA (PERMANOVA; using the metaMDS function in the Vegan package) were performed based on Bray–Curtis dissimilarity matrix. An additional PERMANOVA was used to test for differences in the EMF community composition of saplings, based on identity of the sapling and the identity of the nearest mature tree as well as their interaction. Community compositional differences were visualized using non-metric multidimensional scaling (NMDS).

#### Sapling performance

We used general linear models using a fully factorial design. To account for differences in the sapling height and stem diameter, the following explanatory variables were included as fixed factors: the identity of the sapling, the identity of the nearest mature tree as well as their interaction. Since an observed increase in the number of branches can be simply the result of ontogeny (i.e., larger saplings will have more branches) we included sapling height as a continuous covariate in the model.

The statistical parameters reported are the results of ANOVA omnibus tests for EMF richness, sapling height and stem diameter (data residuals were normally distributed) and log likelihood ratio tests (based on Poisson distribution) for the sapling’s branch number. Figures were generated using R packages ggplot2 (version 3.3).

## Results

### Fungal community richness

ASV’s richness was similar in all samples regardless of the tree species sampled (F_1,65_=0.019, p=0.889) or the tree age (F_1,65_=0.09, p=0.756). Similarly, richness calculated at the agglomerated species level did not significantly differ between the tree species (F_1,63_=1.83, p=0.180), nor between mature and young saplings (F_1,63_=1.76, p=0.189).

### Fungal community composition of mature trees and saplings

The EMF community composition varied among samples of mature trees and saplings of pine and cedar trees (Fig. 1, Fig. S2). Specifically, *Tomentella coerulea* was highly dominant on young pines (51%) and to a lesser extent on young cedars (23%). *Russula densifolia* was associated with trees of all groups but was highly dominant on mature cedar roots (25%) and to a weaker extent on mature pine roots (20%), while being much less abundant on saplings. Mature pines were highly dominated by *Tuber nitidum* which appeared also on many cedar saplings (26%). Other EMF species (clustered together as “Others”, each accounting for less than 1%) were more abundant on mature trees (15% and 12% on mature cedars and pines, respectively) than on young saplings (4% and 6% on cedar and pine saplings, respectively). There was a significant compositional difference among the different age groups (PERMANOVA: Host age; Pseudo F_1,65_=2.60, p=0.029; Fig. 2). However, there was no significant difference among the tree species (PERMANOVA: Host Identity; Pseudo F_1,65_=0.64, p=0.668; Fig. 2) nor was there a significant interaction between the host age and host identity (PERMANOVA: Interaction; F_1,65_=1.81, p=0.050).

**Figure 1.**
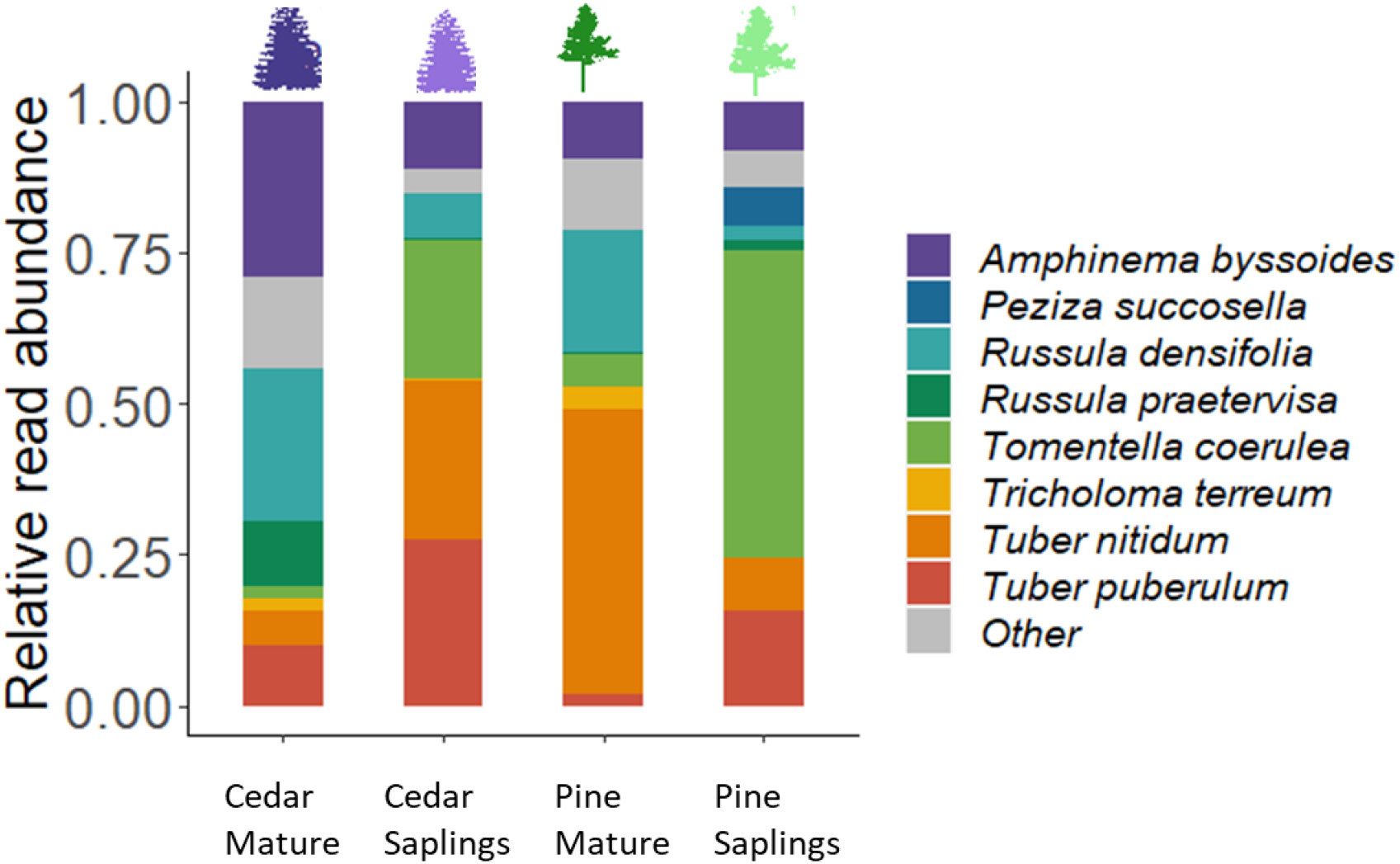
The community composition of the most abundant root-associated ectomycorrhizal fungi, extracted from roots of mature trees and young saplings of *Pinus helepensis* and *Cedrus deodara* in Mt. Meron, Israel.

**Figure 2.**
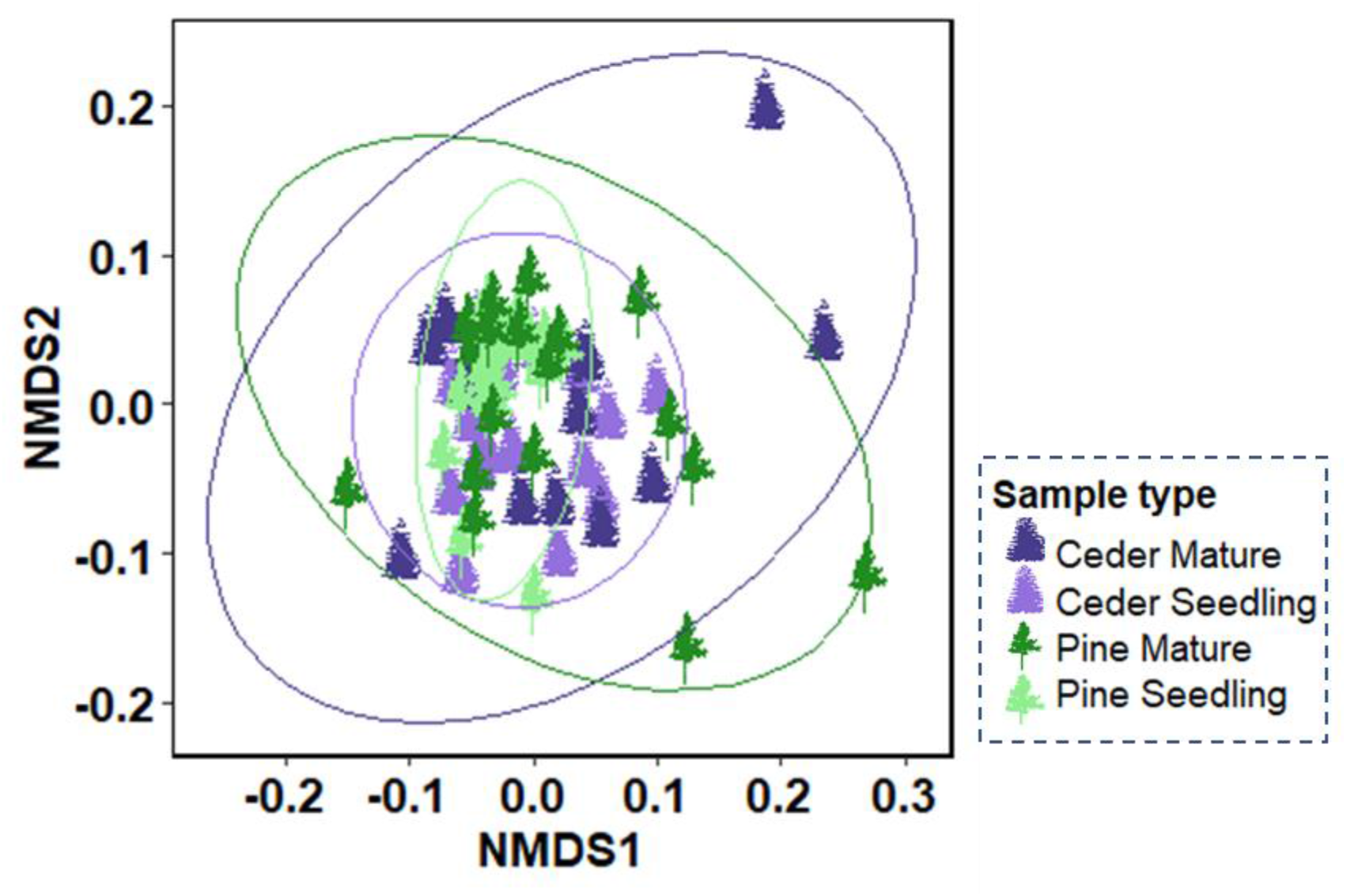
The community composition of root-associated ectomycorrhizal fungi, extracted from roots of mature and young saplings of *Pinus helepensis* and *Cedrus deodara* in Mt. Meron, Israel, and illustrated using non-metric multi-dimensional scaling (nMDS). Colors denote the different sample type (pine/cedar, mature/ seedling). Circles denote ±95 CI.

### The identity of the nearest mature tree affects the EMF community composition of saplings

Examining the different factors contributing to the variation in the fungal communities of saplings (Fig. 3), we found a significant effect of the identity of the nearest mature tree (Nearest mature identity: PERMANOVA; Pseudo F_1,34_=4.00, p=0.001).The overall EMF richness reduced with the distance from mature trees (Pearson’s r= −0.35, p=0.031, Fig. 4), but more importantly, the identity of the nearest mature tree appeared to be shaping the fungal communities of cedar and pine saplings more than the identity of the sapling itself, which appeared to have a non-significant effect on the similarity among saplings. (PERMANOVA: Pseudo F_1,34_=1.35, p=0.18). The interaction between the host identity and the identity of the nearest mature tree was also non-significant (Interaction: PERMANOVA; Pseudo F_1,34_=1.76, p=0.068). This separation among saplings which germinated in proximity to either mature pine or cedar trees, is based mostly on the higher abundance of *Tuber* species on the roots of saplings germinating next to cedar trees and *Tomentella coerulea, Russula densifolia* and *Tuber nitidum* dominating the saplings germinating next to mature pines (regardless of the sapling’s own identity; Fig. S2).

**Figure 3.**
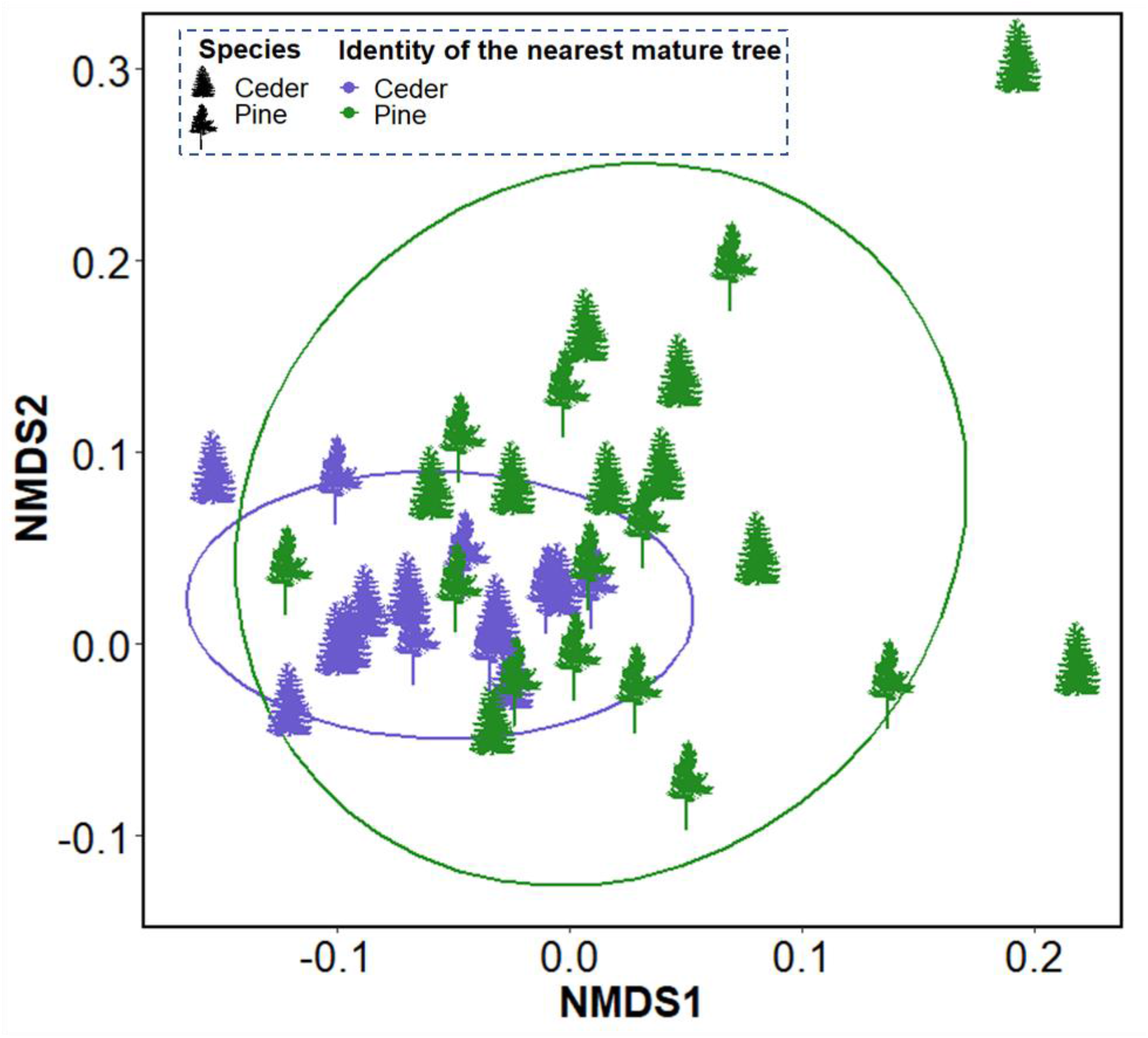
The community composition of root-associated ectomycorrhizal fungi, extracted from young sapling roots of *Pinus helepensis* and *Cedrus deodara* in Mt. Meron, Israel, and illustrated using non-metric multi-dimensional scaling (nMDS). The colors represent the identity of the nearest mature tree from each sampled seedlings. The shapes represent the seedling identity.

**Figure 4.**
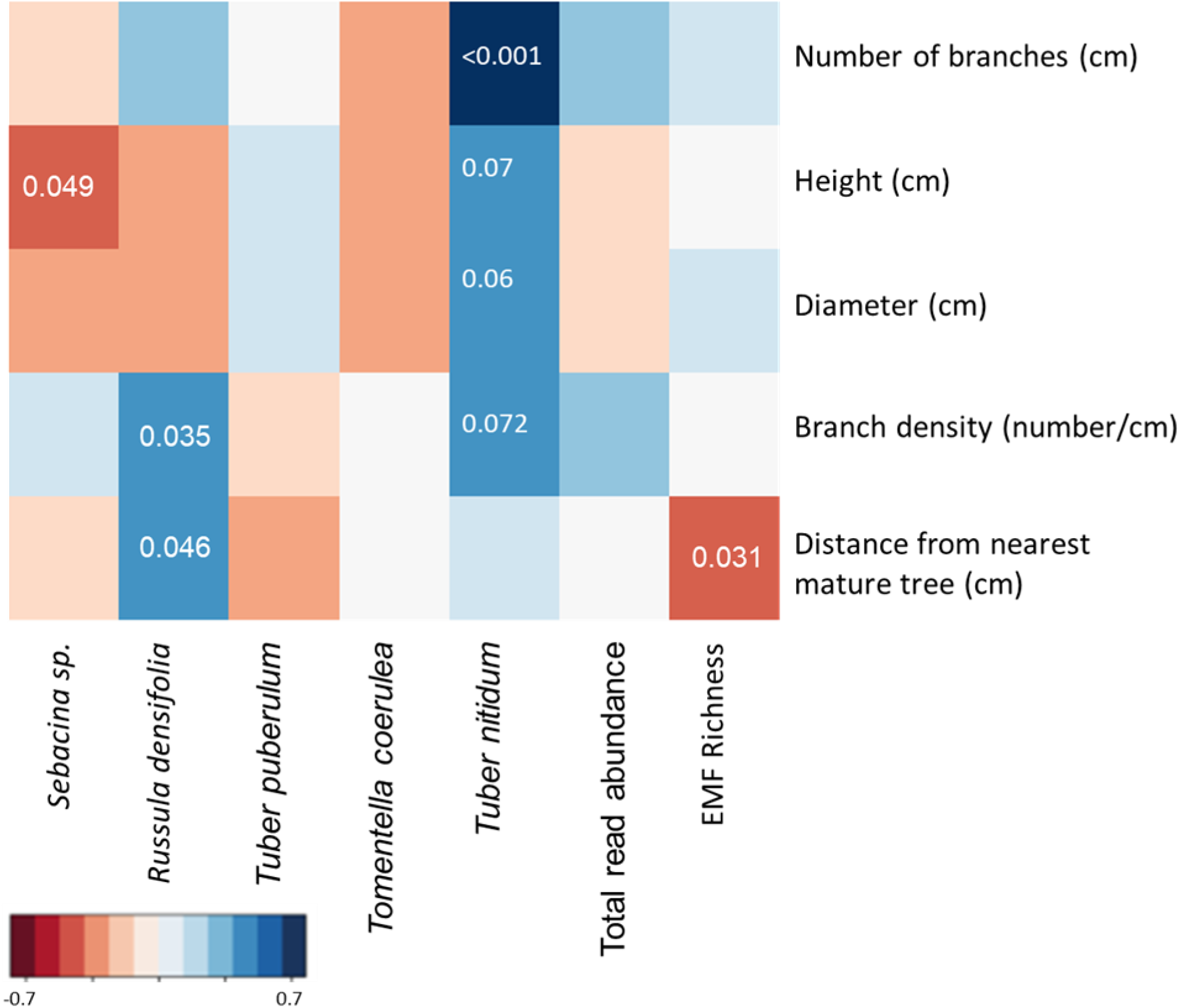
A heatmap presenting Pearson’s correlations between different root-associated ectomycorrhizal fungal species (on the X axis) and different plant performance variables (on the Y axis), extracted from roots of mature trees and young saplings of *Pinus helepensis* and *Cedrus deodara* in Mt. Meron, Israel. Positive correlations appear in blue and negative correlations appear in red. The numbers denote significant/ marginally significant correlations.

### The identity of the nearest mature tree affects sapling performance

We examined the combined effects of the identity of the sapling and the identity of the nearest mature tree on sapling performance. Sapling diameter did not vary among the two tree species (ANOVA: sapling identity; F_1,34_ =3.49, p=0.07) nor was it affected by the identity of the nearest mature tree (ANOVA: sapling identity; F_1,34_=1.04, p=0.314). Pine saplings were slightly taller than cedar saplings (21.05±6.4 cm, ANOVA: sapling identity; F_1,34_=10.81, p=0.002). However, the identity of the nearest mature tree had no effect on sapling height (ANOVA: Nearest mature tree identity; F_1,34_ =0.31, p=0.631). Interestingly, the number of side branches was significantly affected by the identity of the saplings (Log likelihood ratio test: sapling identity; χ^2^_1_=94.25, p<0.01) and the identity of the nearest mature tree (Log likelihood ratio test: Nearest mature tree identity; χ^2^_1_=4.3, p=0.038). Cedar saplings germinating next to mature pines had a significantly higher number of side branches, as apparent from the significant interaction among sapling identity and the identity of the nearest mature tree (Fig. 5; Log likelihood ratio test: interaction; χ^2^_1_=13.38, p<0.01) and this pattern remained when including the sapling height as a covariate (Table S1).

**Figure 5.**
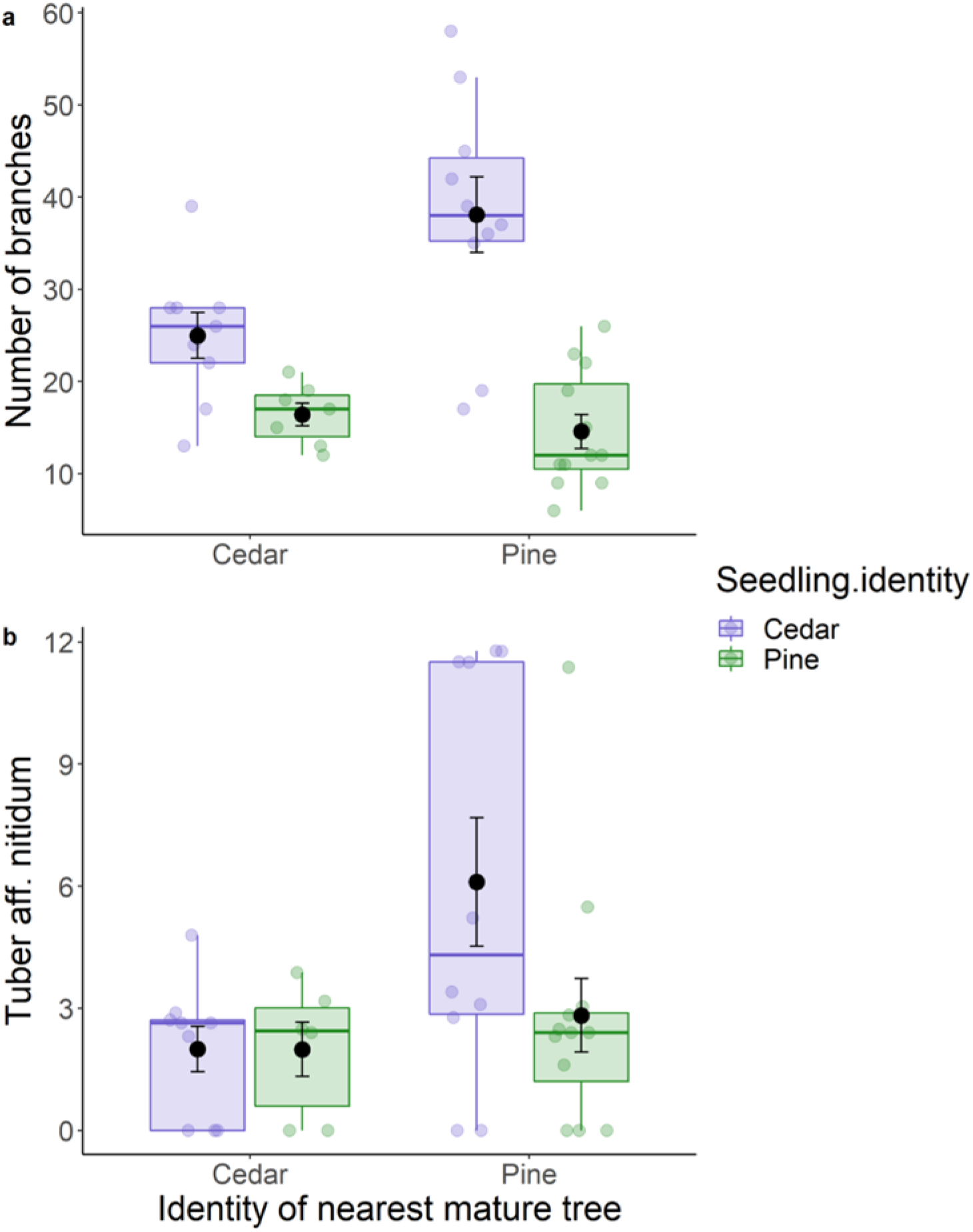
(a) The number of branches, and (b) the relative abundance of *Tuber nitidum* as a function of the identity of the nearest mature tree and the identity of the sapling itself (green=pine, purple=cedar). Data are from roots of young saplings of *Pinus helepensis* and *Cedrus deodara* in Mt. Meron, Israel

When examining the simple correlations between the relative abundance of the most abundant fungal taxa and the various measures of sapling performance (Fig. 4), the number of branches (Pearson’s r= 0.63, p<0.001, Fig. 3) and to some extent sapling stem diameter (Pearson’s r= 0.31, p=0.06) shoot length (Pearson’s r= 0.30, p=0.07) and branch density (Pearson’s r= 0.29, p=0.072), were all positively correlated with the relative abundance of *Tuber nitidium*. Similar to the pattern of branch density, cedars germinating next to mature pines had a higher relative abundance of *Tuber nitidum* on their roots (Fig. 5).

## Discussion

EMF community richness and composition is expected to vary among host trees and to change along the tree’s ontogeny. In this study we examined the EMF community and how two co-habiting trees (*Pinus halepensis* and *Cedrus deodara*), naturally regenerating in a post-fire site, change with host age. We predicted that young saplings would have a unique mycorrhizal community distinguishing each host species. However, we found that the composition of the saplings’ EMF community was more affected by the identity of the nearest mature tree than by the identity of the sapling. These differences in the EMF community composition were correlated with sapling morphology, but not size. Furthermore, we found that the differences between the two species were mostly in the relative abundance of the species in the community and not in species composition.

### Pines and Cedars have similar EMF communities

Both pines and cedars are closely related gymnosperms, belonging to the Pinaceae (but are quite separated within this clade) which might explain the similarity in their mycorrhizal communities (Ishida et al., 2007, Glassman et al., 2017, Nguyen et al., 2016; Fig. 1). However, we expected mature trees to be more similar in their EMF community composition in comparison to young saplings, since specific mycorrhizal fungi associated with seedlings are expected to disappear during succession (Thompson et al., 2022, Twieg et al., 2007). Nevertheless, we could not detect such differences, possibly because high host specificity is only apparent in the early seedling stages of these pines and cedars and became less apparent as the trees aged from seedling to saplings at the time our sampling took place.

### Mode of inoculation explains lack of difference in EMF community composition in saplings

Another possible origin of the small differences in EMF community composition between the saplings of the two species could be related to their mode of inoculation. A large share of the current knowledge regarding the EMF community of young tree seedlings originates from bioassay-based studies (Taylor and Bruns, 1999, Horton and Bruns, 1998, Baar et al., 1999). Under bioassay settings, colonization of roots is almost entirely restricted to spore-based inoculum. This restriction might give an advantage to members of the *Suilloids*, which are known to have hardy spores that germinate quickly (Hayward et al., 2015). When seedlings germinate under natural conditions this advantage might be less prominent, depending on the availability of different inoculum sources in the soil. In a study conducted after a stand replacing fire, Glassman *et al*., (2016) found that the EMF community on seedling roots was similar and highly dominated by *Suilloids* both in a bioassay and in seedlings grown in the field. This lack of difference could originate from the fact that a stand replacing fire kills off all the hosts, leaving behind only spore-based inoculum sources. Under less intense disturbances, hyphae are expected to withstand, enabling other species that have a more hyphae-based strategy, to colonize the seedling roots. In such cases that adult tree identity and location can affect the regenerating EMF community. In concordance with this assertion, Cline et al., (2005) showed that the EMF communities of seedlings planted near trees were more similar to those of mature trees, while seedlings far from trees were more similar to greenhouse bioassay seedlings. In our study, the occurrence of living mature trees in the vicinity of the saplings could have resulted in hyphae based inoculation and reduced differences between the two species.

### Inoculation from spores compared to hyphae

In many post-disturbance systems (e.g., post-fire) the mature host trees are lacking and the main inoculum source for regeneration are spores (Glassman et al., 2016). However, in our study, several mature trees remained, surrounding the area where young pines and cedars were establishing (Fig. S1). Indeed, when examining the similarity among the young cedars and pines in our study, it appeared that the identity of the nearest mature tree had a significant effect on the EMF communities of these saplings (Fig. 3). There are two main pathways by which the presence of a mature tree can affect the EMF community composition of the young saplings: 1) by increasing the density of fungal spores deposited into the soil spore bank (Peay et al., 2012) and 2) by increasing the abundance of fungal hyphae allowing the young trees to “tap” into the existing mycelial network (Teste and Simard, 2008). Our study design does not allow us to separate these two mechanisms by which mature trees can facilitate EMF colonization of young trees. However, members of the Pinaceae are known to be associated with specific fungal taxa such as *Suillus* and *Rhizopogon* dominating the soil spore banks in many post-disturbance habitats (Glassman et al., 2016, Kjøller and Bruns, 2003). Nevertheless, we did not observe these taxa to be highly dominant in our study, further suggesting that our young saplings did not depend on colonization from spores as the main source of fungal colonization.

In our study, EMF richness tended to reduce as the distance from the nearest mature tree increased. The distance from a mature tree has been previously shown to have a non-linear effect on the performance of young trees, and the optimal distance is such that the mature tree canopy does not shade the young seedling but does increase the EMF richness in the seedling surrounding (Dickie et al., 2005).

### The association with specific fungi affects sapling structure

The proximity to a specific mature host tree was correlated with the EMF community composition but not associated with the size of the cedar and pine saplings (i.e., shoot length and stem diameter). However, it was significantly correlated with the number of side branches, specifically, cedars germinating next to mature pine trees had a higher number of branches (regardless of their size, see Table S1). Several studies have shown that the interaction with specific fungal taxa can influence the performance of young seedlings (Bennett and Bever, 2007, Bent et al., 2011). Sapling structure (number of branches and branch density) was positively correlated with the abundance of specific fungal taxa i.e., *Tuber nitidum*. Interestingly, cedar saplings germinating in high proximity to mature pine trees had higher association with this specific fungal taxon (Fig. 5), which was relatively abundant on mature pines (Fig. 1). This result emphasizes the importance of fungal identity in comparison to fungal richness in determining plant development (Policelli et al., 2019). It therefore seems that the identity of both the mature tree and the fungi it associates with, might affect the community composition of young seedlings and their branching pattern, respectively.

### Lack of differences in fungal richness between mature trees and saplings

Unlike differences in relative abundance, we could not detect differences in richness between the four plant groups. According to the “island biogeography” theory, species number is expected to increase with increasing “island” size (MacArthur and Wilson 1967). We can assume that the size of the tree “island” is affected by its age, where larger trees have larger and more complex rooting systems which are able to support a higher number of EMF species. Glassman et al., (2017) showed that in a subalpine system, trees served as “islands” for ectomycorrhizal fungi, with EMF richness increasing as a function of the “island” size and age. In our study, fungal richness was similar for mature and young saplings (for both ASV and EMF richness). This difference might be related to the density of the trees in the two field sites. Unlike the sparsely alpine basin sampled in Glassman et al. (2017), the saplings in our study germinated close to mature trees, possibly enabling them to be inoculated by existing hyphae present on the roots of the mature trees. Although we could not detect an increase in fungal richness with age, the EMF communities of mature trees had different relative abundances than those of young saplings. The most apparent difference in the communities of saplings and mature trees was the increase in the relative abundance of *Russula denshfolia* on mature tree roots (Fig. 1). This fungal species, characterizing mature forest trees, was shown before to become more dominant as the forest matured during forest succession in a study of the EMF communities of Douglas-fir and paper birch along a chronosequence of forest development after stand-replacing disturbance (Twieg et al. 2007).

### Sampling methods and its effect on richness results

The sampling scheme used in our study might have also contributed to the lack of difference in EMF richness between mature trees and saplings (Chase et al., 2018). Specifically, we dug out the saplings and calculated the richness of their entire root system. In mature trees however, richness was calculated after sampling only a very small fraction of the roots. This limitation was further exacerbated by the fact that we could not clump together the five roots sampled around each tree because of the large uncertainty regarding the root origins. This is not the first time that DNA verification of mature roots sampled under field conditions shows such low correct identification rates. Rog et al., (2020) reported similar misclassification of mature roots while sampling mature trees in a temperate mixed forest, and even higher rates of misclassification were observed while sampling mature roots in a dense and dry mixed Mediterranean forest (unpublished data). Overall, such misclassification could be expected, especially in dry and dense forests with shallow soil layers where many roots are tangled in the topsoil layer, highlighting the importance of molecular identification of individual roots.

### Summary and importance

To comprehensively disentangle the intrinsic (i.e., host identity) and extrinsic factors (i.e., identity of the surrounding trees) shaping seedlings mycorrhizal communities, a more comprehensive study across a larger geographical scale, involving a higher number of hosts, is needed. However, this unique case-study provided us with an opportunity to test how the spatial distribution of surviving trees in a post-disturbance habitat affected both the EMF communities and the performance of the regenerating saplings. Understanding the various factors governing forest regeneration (both aboveground and below) is essential when managing both natural and silviculture (Liira et al., 2011) habitats. With mature trees becoming highly vulnerable, the importance of seedling establishment is highlighted and with it the need to understand how this process is influenced by their mutualistic interactions.

## Acknowledgments

We wish to thank Shahar Yirmiahu, Yair Zach, Amal Qassem, Yuval Tauber, Amit Berlinsky and Oren Lugasi for their assistance during sampling and to Sophie Obersteiner for her help with molecular verification of mature roots. We wish to thank the National Park Authority for permitting us to work within the national park. We are grateful to Yvonne Lipman for the English editing. The data regarding the tree species was gathered using the TRY initiative on plant traits (http://www.try-db.org). The TRY initiative and database are hosted, developed and maintained by J. Kattge and G. Bönisch (Max Planck Institute for Biogeochemistry, Jena, Germany). TRY is currently supported by DIVERSITAS/Future Earth and the German Centre for Integrative Biodiversity Research (iDiv) Halle-Jena-Leipzig.

## Author Contributions

SLL, HS, and MA conceived and designed the experiment. MA, IR, and LH collected the data. MA, IR, and SLL analyzed the data. SLL wrote the paper with MA, TK, and HS, and all authors contributed substantially to revisions.

## Data availability statement

Sequences were submitted to the National Center for Biotechnology Information Sequence Read Archive with the accession codes: Bioproject PRJNA777920.

