## Supplementary information for "Tell me who your neighbors are: The role of spatial location and tree species identity in determining the ectomycorrhizal community composition of saplings and mature trees in a mixed conifer forest"

**Supplementary material**

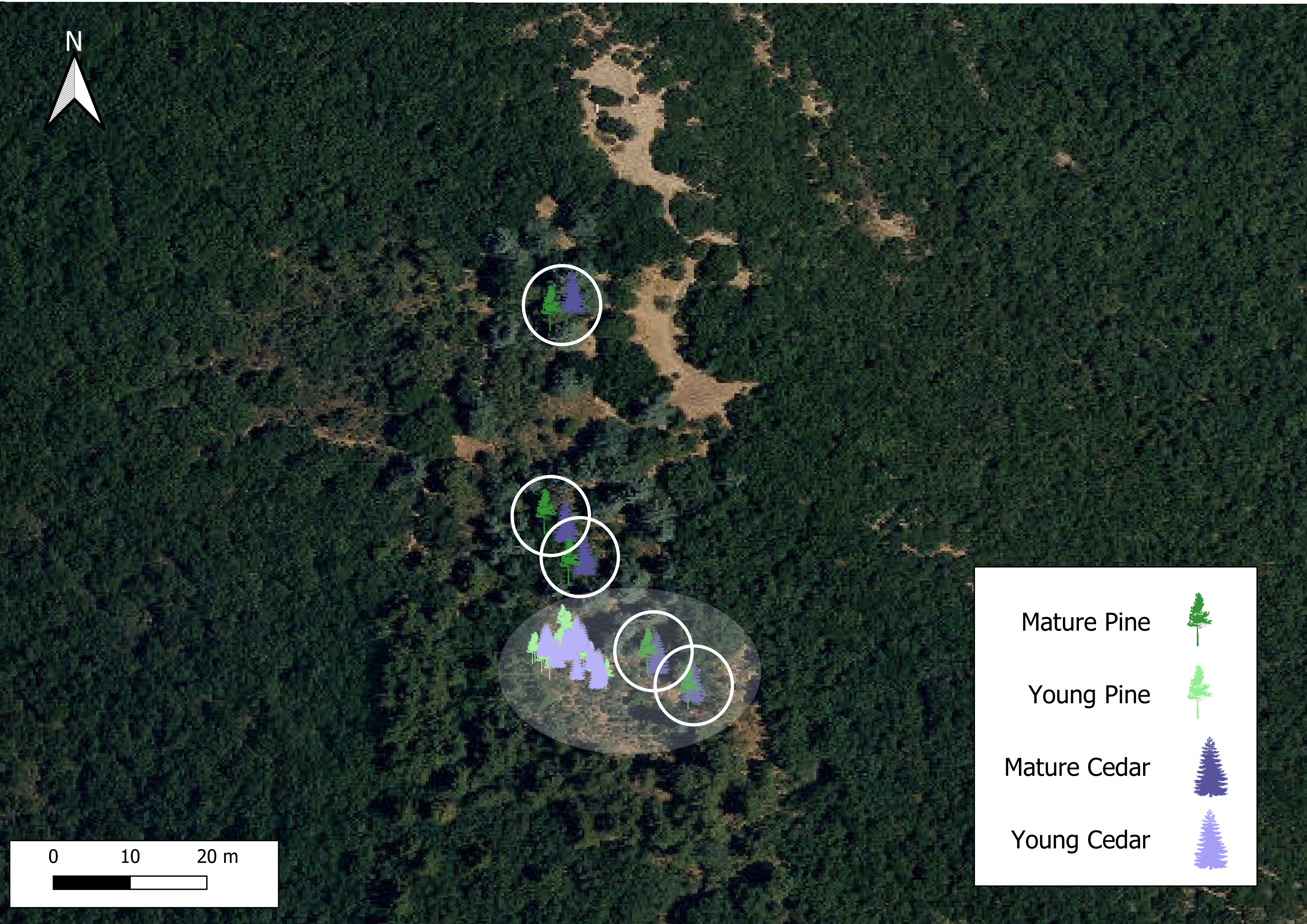

Figure S1. A map representing the locations of the sampled mature trees (five pairs denoted in circles) and young saplings (37 in total) of *Pinus helepensis* and *Cedrus deodara* in Mt. Meron, Israel.

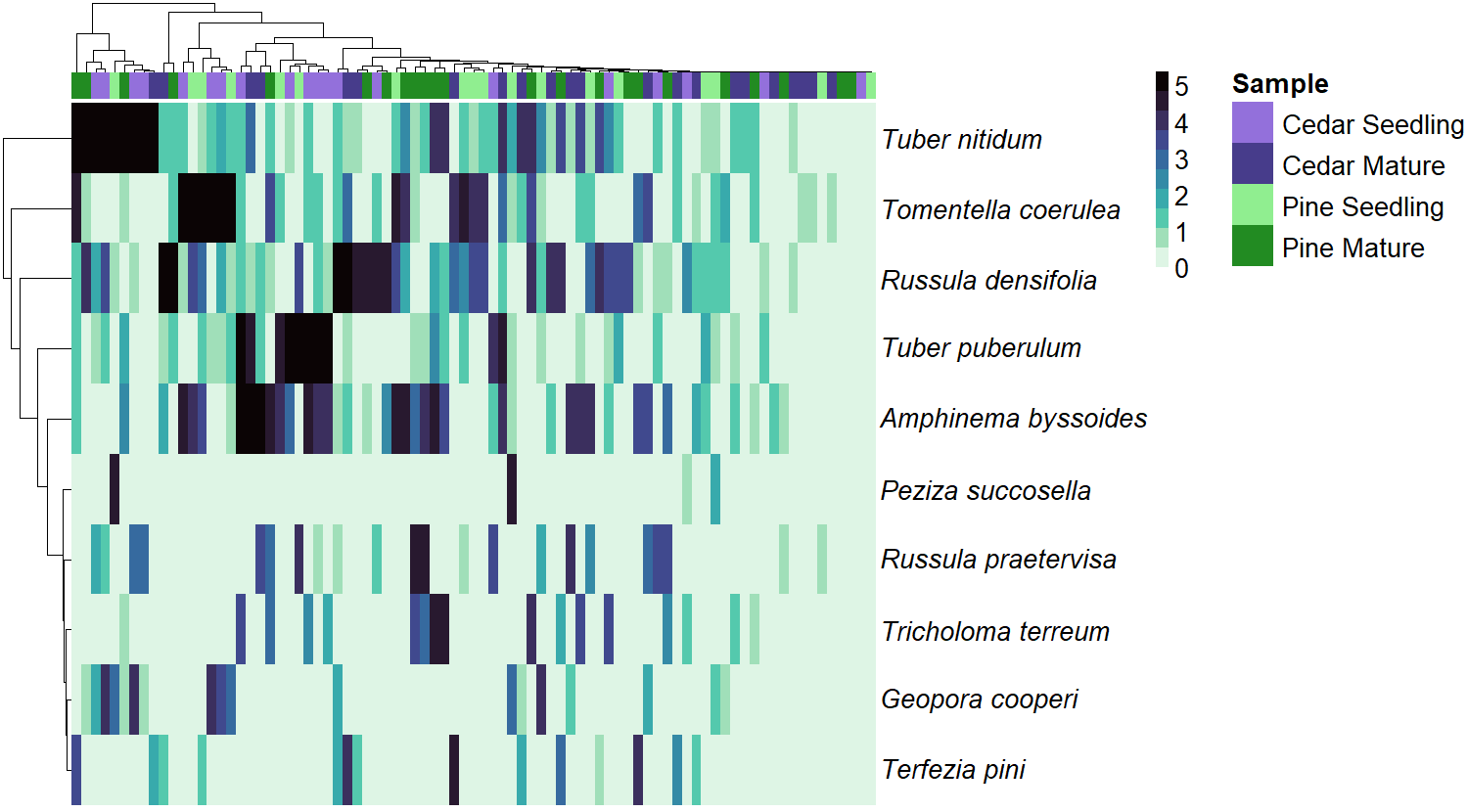

Figure S2. A heatmap representing the log10 relative abundance of root-associated ectomycorrhizal fungi, extracted from roots of mature trees and young saplings of *Pinus helepensis* and *Cedrus deodara* in Mt. Meron, Israel.
